# Environmental DNA reflects common haplotypic variation

**DOI:** 10.1101/2022.02.26.481856

**Authors:** Clare I M Adams, Christopher Hepburn, Gert-Jan Jeunen, Hugh Cross, Helen R Taylor, Neil J Gemmell, Michael Bunce, Michael Knapp

## Abstract

Analysis of environmental DNA (eDNA) has gained widespread usage for taxonomically based biodiversity assessment. While interest in applying non-invasive eDNA monitoring for population genetic assessments has grown, its usage in this sphere remains limited. One barrier to uptake is that the effectiveness of eDNA detection below the species level remains to be determined for multiple species and environments. Here, we test the utility of this emergent technology in a population genetic framework using eDNA samples derived from water along New Zealand’s South Island (Otago Coast: n=9; Kaikōura: n=7) and DNA obtained from tissue samples (n=76) of individual blackfoot pāua (*Haliotis iris)* sampled from New Zealand’s Otago coast. We recovered four mitochondrial haplotypes from eDNA versus six from the tissue samples collected. Three common haplotypes were recovered with both eDNA and tissue samples, while only one out of three rare haplotypes – represented in tissue samples by one individual each – was recovered with our eDNA methods. We demonstrate that eDNA monitoring is an effective tool for recovering common genetic diversity from pāua, although rare (< 5%) haplotypes are seldom recovered. Our results show the potential of eDNA to identify population-level haplotypes for gastropods in the marine environment identification below the species level and for studying the population genetic diversity of gastropods. This work supports eDNA methods as effective, non-invasive tools for genetic monitoring. Non-invasive eDNA sampling could minimize target organism stress and human interaction enabling population genetic research for hard-to-sample, delicate, or sensitive species.

## 1. Introduction

Environmental DNA technologies are evolving rapidly. An emerging trend is the transition from genetic markers that identify species to those that evaluate population-level polymorphisms (e.g. Parsons et al. 2018, Stepien et al. 2019) to enable population genetic variability to be discerned using eDNA (Sigsgaard et al., 2020) Obtaining population genetic data from eDNA could potentially help estimate population size, inbreeding, gene flow, and population viability (Adams et al., 2019).

Population-level genetic information in the form of haplotypes has been obtained with eDNA methods for a variety of taxa, including dolphins, fish, and bivalves (Marshall & Stepien, 2020; Parsons et al., 2018; Sigsgaard et al., 2016; Stepien et al., 2019; Tsuji et al., 2020). More recently, biotinylated baits have been used to capture mitochondrial and nuclear fish SNPs from water samples (Jensen et al., 2020). However, eDNA population genetic results are only useful if we can be confident that eDNA methods are capturing a robust estimate of the genetic variation present in the target population. Field experiments that compare the results from traditional population-genetic sampling methodologies and analyses to those using eDNA based population genetic estimates are required. Here, we seek to determine if non-invasive eDNA samples targeted to capture the population genetic diversity of a marine gastropod will produce the same results as traditional population-genetic surveys using invasive tissue sampling.

The blackfoot pāua (*Haliotis iris*) is an abalone species endemic to Aotearoa New Zealand. The blackfoot pāua fishery had a $15 million NZD export value in 2020 (Seafood New Zealand, 2020), and was valued at $80 million NZD in exports at its 2001 peak (The Pāua Industry Council, 2021). Importantly, blackfoot pāua is also a cultural keystone species for Māori and are considered a tāonga (treasured species) for Ngāi Tahu, a major iwi (tribe) in the South Island of New Zealand (Garibaldi & Turner, 2004; McCarthy et al., 2014; Turner et al., 2013). Traditionally, Māori exerted kaitiakitanga (guardianship) over the marine environment for the harvest of kaimoana (seafood) using mātauranga (indigenous knowledge) and tikanga (practice), but European colonialism and centralized management have since undermined efforts for sustainable harvest (Bess 2001, McCarthy et al. 2014, Bennett-Jones et al. 2021). Harvesting of blackfoot pāua is managed using a variety of tools including total allowable catch and minimum legal size for commercial fisheries, and daily bag limits and minimum size for recreational fishing and authorisations for traditional fishers guided by local mātauranga (Gnanalingam et al., 2021). However, like most wild abalone populations, blackfoot pāua populations are still declining (Cook, 2016). Analyses using microsatellite and mitochondrial markers have suggested that blackfoot pāua have relatively high genetic diversity within populations, and weak, but significant, structure between populations (Will, 2009; Will et al., 2011, 2015).

The cultural importance of pāua, the broadly decreasing populations of abalone worldwide (Cook, 2016), and the high genetic diversity of blackfoot pāua make them a practical choice for testing how effective eDNA is at obtaining haplotypic genetic diversity (Cook, 2016). In addition to this, blackfoot pāua individuals are easy to capture and have an already-sequenced mitogenome, making eDNA and tissue comparison feasible (Guo et al., 2019). While other studies have compared eDNA-obtained haplotypes to previously obtained population genetic data, few population-level eDNA studies have taken both tissue and eDNA at the same time in the field (Marshall & Stepien, 2019; Parsons et al., 2018; Székely et al., 2021). Therefore, we ask: (1) Can we obtain representative blackfoot pāua eDNA haplotypes from a field setting? (2) Does eDNA reflect tissue-based haplotypic diversity from a field setting? (3) Is the assay applicable to other locations? This work helps further solidify eDNA methods for population genetic information and for use in fisheries management. We also make recommendations on sampling strategy for identifying rare haplotypes based on our results.

## 2. Materials and Methods

### 2.1 Sample collection and DNA extraction

To compare eDNA and tissue samples, we took eDNA water samples and then immediately harvested blackfoot pāua (*Haliotis iris*) individuals around a rock off the coast of Warrington, Otago on 16 April 2019 (location withheld due to poaching concerns) (Figure 1). Because blackfoot pāua are patchily distributed, our water collection occurred around a single blackfoot pāua-populated rock (Figure 2). For eDNA samples, nine 2L samples of water were collected around the pāua-populated rock in acid-washed, glass bottles. These bottles were taken down five meters, opened, rinsed, and purged with air from a SCUBA regulator. Once all water samples had been collected, we collected 76 adult blackfoot pāua individuals at random from around that same rock (MPI Special Permit to the University of Otago (644)). Animals were transported live back to the University of Otago’s Portobello Marine Lab (PML) for spawning to support local blackfoot pāua fishery restoration.

**Figure 1.**
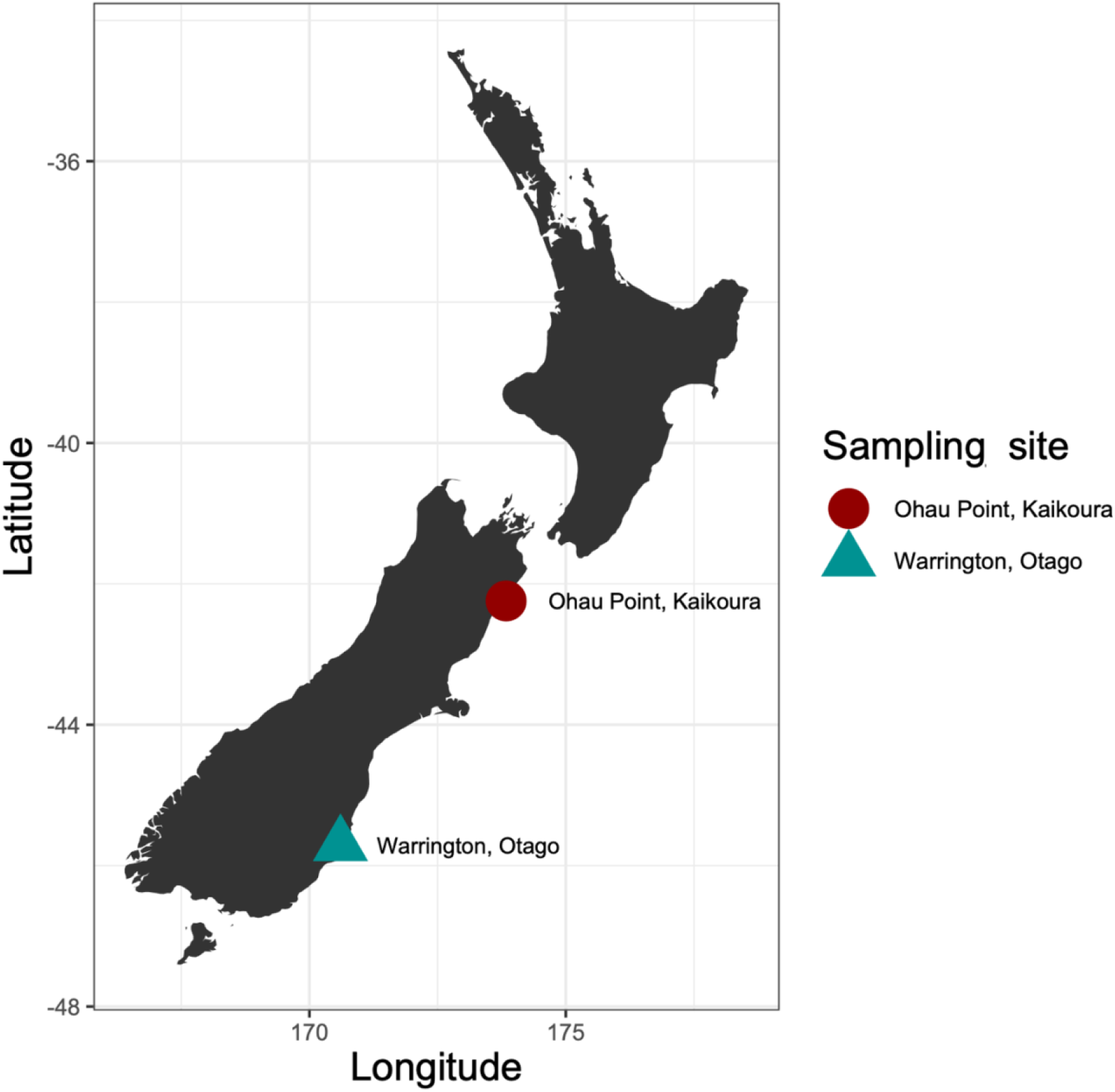
Map of sampling sites Ohau and Warrington. The distance between them is approximately 450 km.

**Figure 2.**
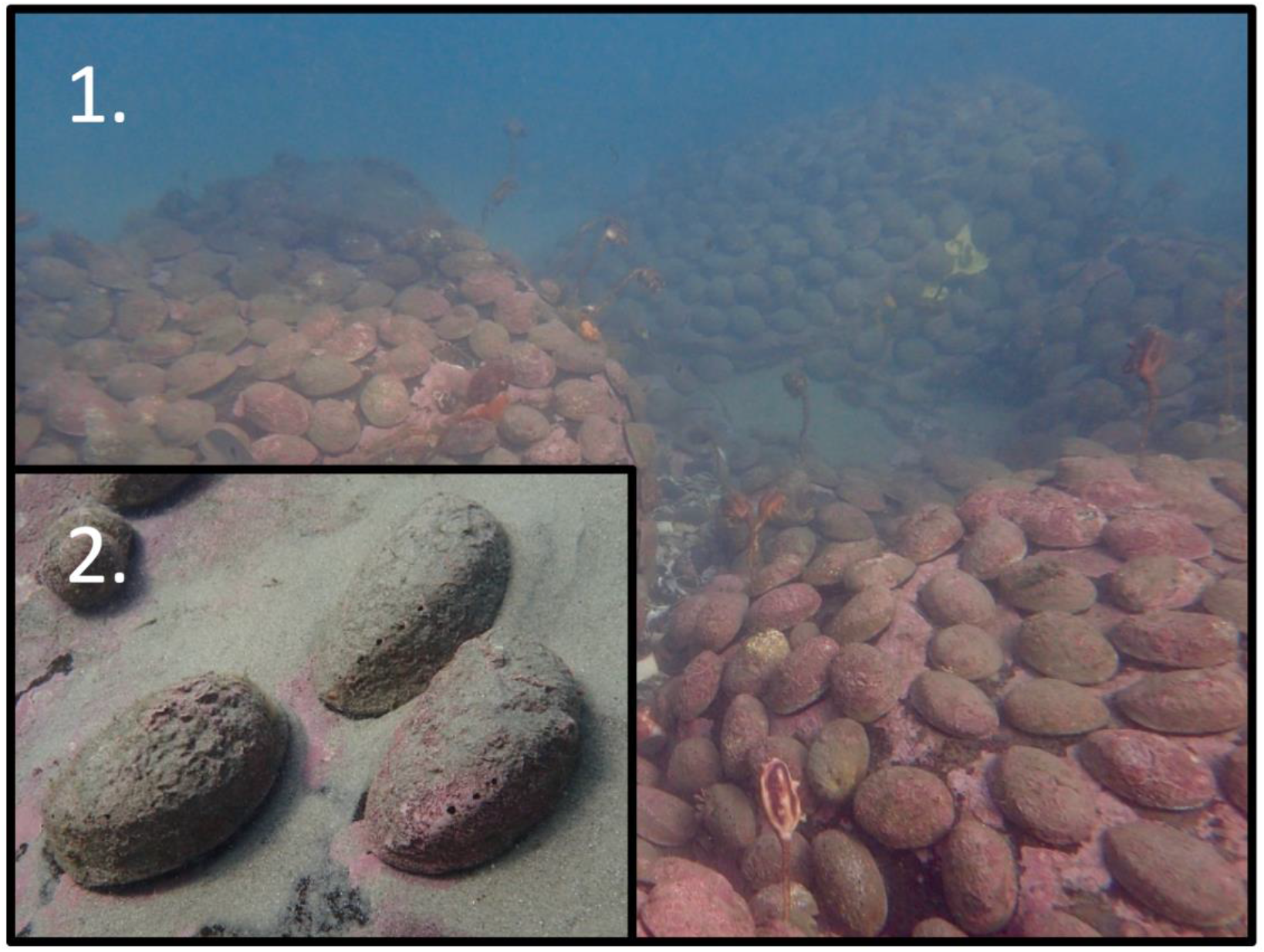
An image representative of pāua beds off the coast of Warrington, Otago, New Zealand. (1) Blackfoot pāua (*Haliotis iris*) beds spread across rocks in a patchily distributed population. (2) Three individual blackfoot pāua covered in sand. Photo credit: Dr. Matthew Desmond.

To determine if blackfoot pāua haplotypic variation could be recovered from eDNA at other sites, we opportunistically collected seven 500mL eDNA samples from the shore near Ohau Point, Kaikōura on 13 February 2020 (Figure 1). This location had blackfoot pāua shells scattered across the rocky beach and this site was known as prime pāua habitat prior to the 2016 Kaikōura earthquake (Fisheries New Zealand & Kaikōura Marine Guardians, 2018).

We filtered eDNA samples immediately at PML (Warrington) or in the field (Ohau Point) with Sterivex (0.22μm) columns (Millipore) (Spens et al., 2017). For Warrington, three separate 500mL replicates were taken per sample, and negatives were taken before and after sampling. This resulted in nine samples with three 500mL replicates each, totalling 1.5L filtered for each sample. Three mL of Longmire’s buffer was added to each Sterivex filter to preserve samples (Figure 3). For Ohau Point samples, each of the seven 500mL eDNA samples were filtered on-site according to Spens et al., (2017). Three mL of Longmire’s buffer was added to preserve eDNA in the Sterivex filter during transport back to Dunedin, Otago. Five hundred mL of ultrapure DNase/RNase-free water (Invitrogen) was filtered before and after sampling for use as negative field controls at both locations. All eDNA samples were extracted within a week using Qiagen Blood and Tissue Kits in the Otago University PCR-free eDNA laboratory (Spens et al., 2017). All laboratory equipment was bleached prior to use and a negative extraction was added during extraction to ensure clean laboratory practice.

**Figure 3.**
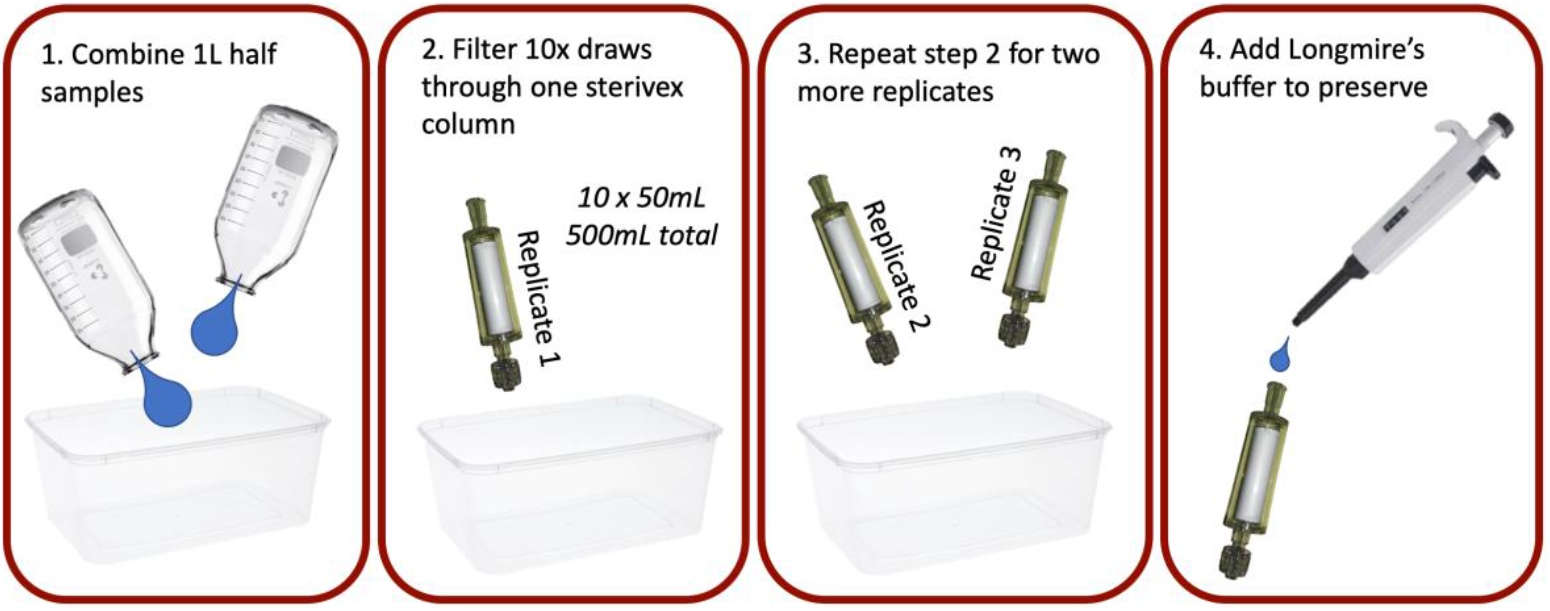
Graphical image of how samples were filtered. For each sample, (1) each 1L half sample was combined for a 2L total sample. (2) Ten draws of a sterivex filter through a 50mL syringe (not pictured) resulted in a 500mL replicate. (3) Three replicates were taken per sample. (4) Longmire’s buffer was added to each sterivex column for preservation.

Live blackfoot pāua (n=76) from Warrington were housed at PML for epipodial (tentacles on the mantle) tissue sampling. Bench surfaces were cleaned with 10% bleach before work. Between three and seven epipodia were plucked from each individual and placed in a 2mL tube with 1mL of 95% ethanol. Tweezers were washed with 10% bleach and 95% ethanol between individuals. Pāua individuals were then returned to their aquaria at PML for an experimental breeding program and teaching laboratory use. Pāua epipodial samples were taken back to the PCR DNA laboratory (PCR allowed) and extracted within a week. We used the Qiagen Blood and Tissue Kit according to manufacturer’s instructions for these tissue samples, with the addition of an extended incubation overnight in ATL lysis buffer. Negative extraction controls containing no tissue were extracted alongside pāua tissue extractions and carried through library preparation every eight to 16 samples. Laboratory equipment and bench surfaces were cleaned with 10% bleach and rinsed with dH2O before work.

### 2.2 Pāua tissue library preparation and analysis

DNA extracts from tissue samples were used to construct partial mtDNA genotypes for all (n=76) individuals. Novel long-range primers were combined with primers from Gou et al., (2019) to amplify blackfoot pāua mtDNA (Table 1). Two long-range PCRs amplified pāua fragments from 812bp - 13056bp (Table 1) in a 30μL reaction volume containing 1x KAPA LongRange Buffer (without Mg^2+^), 1.75mM MgCl2, 0.3mM dNTPs, 0.5μM each primer, 2μL of template, and 1.5 U KAPA LongRange HotStart DNA Polymerase. No-template controls were included in this and the following PCRs. Thermal cycling conditions had an initial denature at 94°C with 35 cycles of a 94°C denature for 30 seconds, 55°C annealing at 30 seconds, and 68°C extension for 10 minutes. This was followed by a final extension at 68°C for 20 minutes. Primer pairs designed by Gou et al., (2019) – HI8, HI9, and HI10 (Table 1) – were amplified in a different 25μL reaction using 1x KAPA Taq Buffer, 0.6 mM dNTPs, 0.4μM each primer, 0.5U KAPA Taq DNA Polymerase, and 1μL of template DNA. Cycling conditions were an initial denaturation at 95°C for 5 minutes, followed by 30 cycles at 95°C for 30 seconds, 58°C for 30 seconds, 72°C for 3 minutes, and then a final extension at 72°C for 10 minutes, with the exception of HI9, which annealed at a temperature of 51°C. Note that there are gaps between base pairs (bp) 13,056-13,073 and 15,576-16,646 and 1-812 (Table 1). These regions could not successfully be amplified due to either little to no amplification or multiple bands.

**Table 1.**
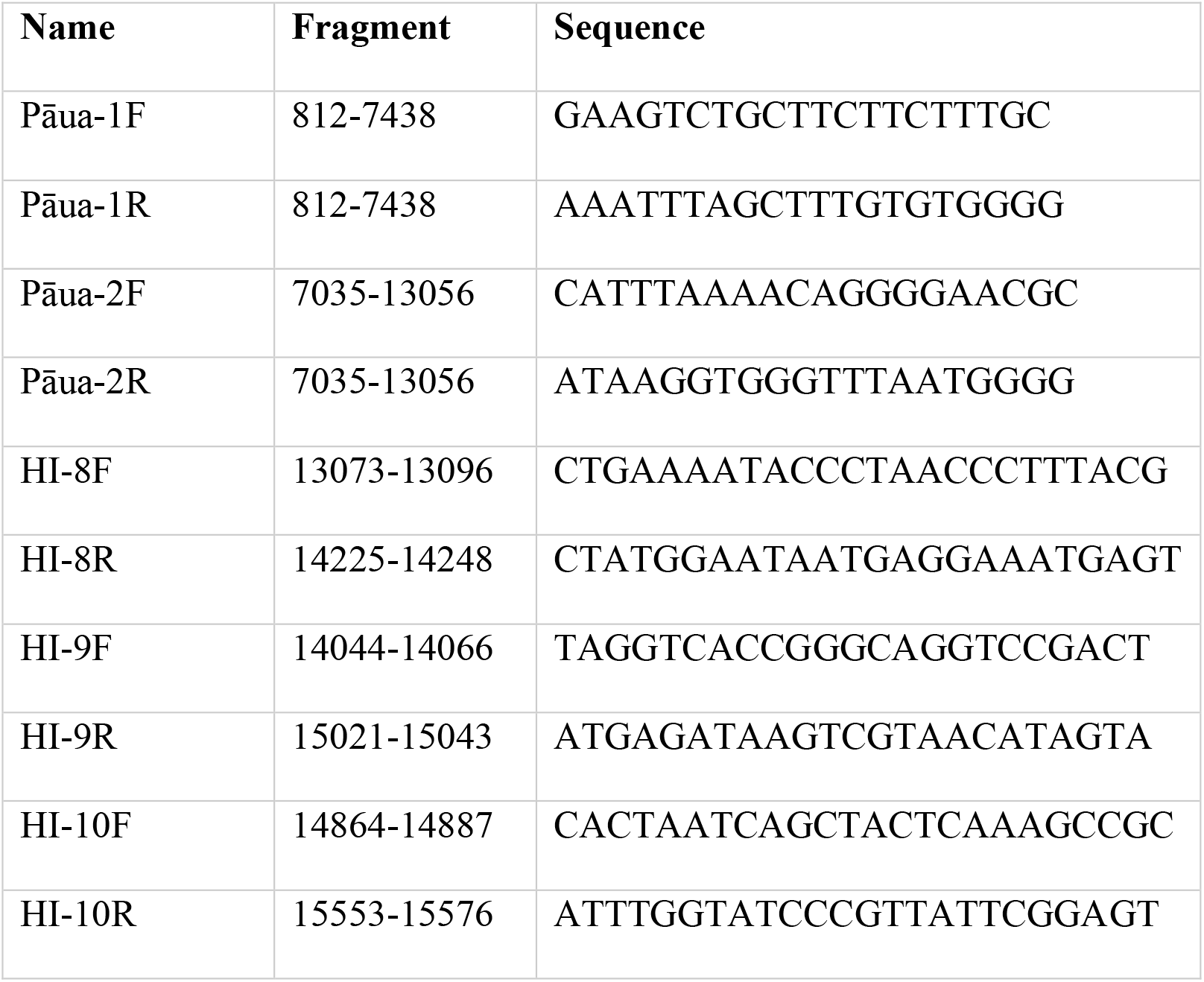
Primers used for amplifying blackfoot pāua (*Haliotis iris*) tissue samples.

We prepared double-stranded DNA libraries for pāua tissue samples, including negatives (Kircher et al., 2012). Sonication was performed with a Covaris S220 focused-ultrasonicator at 15 seconds on and 90 seconds off for eight cycles at 4°C. We confirmed successful shearing of products to less than 100 bp with gel electrophoresis. Blunt end repair and adapter ligation was performed according to Knapp et al., (2012) with the exception of clean-up using AMPure XP beads instead of a MinElute Kit (Qiagen). Amplified products were quantified with Quant-iT(tm) PicoGreen dsDNA Assay Kit on a VICTOR Multilabel Plate Reader (PerkinElmer) and pooled in equimolar ratios. We submitted the pooled library to Otago Genomics for paired-end 2×300 sequencing.

Once reads were obtained, we ran the demultiplexed samples through the Eager 2.0.7 pipeline using NEXTFLOW v19.01.0 with modifications for modern mitochondrial data and using the *H. iris* mtDNA genome as a reference (NC_031361.1) (Guo et al., 2019; Yates et al., 2021). We extracted the consensus sequences for each sample and aligned them using Geneious Prime 2020.1.1 (https://www.geneious.com) (Table 2). Seventy-one of the seventy-six blackfoot pāua samples were successfully sequenced in the desired amplicon range and these were used for downstream analyses and comparisons with eDNA amplicons (Table 2).

**Table 2.** Mitochondrial reads per sample, coverage, and read depth from blackfoot pāua tissue from Warrington, Otago when mapped to the blackfoot pāua genome (NC_031361.1).

|  | Reads per Sample |  |  | Coverage |  |  | Read Depth |  |  |
| --- | --- | --- | --- | --- | --- | --- | --- | --- | --- |
|  | Range | Average | St Dev | Range | Average | St Dev | Range | Average | St Dev |
| *Haliotis iris* | 1,333 - 59,817 | 21766.37 | 10761.74 | 55.28 - 86.33 | 85.50 | 3.97 | 23.60 - 908.172 | 334.20 | 441.58 |

### 2.3 eDNA Primer development

Previously published primers amplified across abalone species (Will, 2009) and long-range primers may have been too long for low-concentration, degraded eDNA (Barnes et al., 2020; Moushomi et al., 2019). Hence, novel eDNA primers were developed that included a region between mitochondrial proteins for species-specificity. Based on our sequences from blackfoot pāua tissue, specific eDNA primers were designed in Geneious Prime 2020.1.1 (https://www.geneious.com) using Primer3 across the ATP8 and ATP6 regions (bp 5,875 – 6,183 in KU310895) (Table 3). Aligned consensus sequences from each individual were compared to determine the presence of three common haplotypes, represented by more than two individuals, and three rare haplotypes, representing one individual each.

**Table 3.**
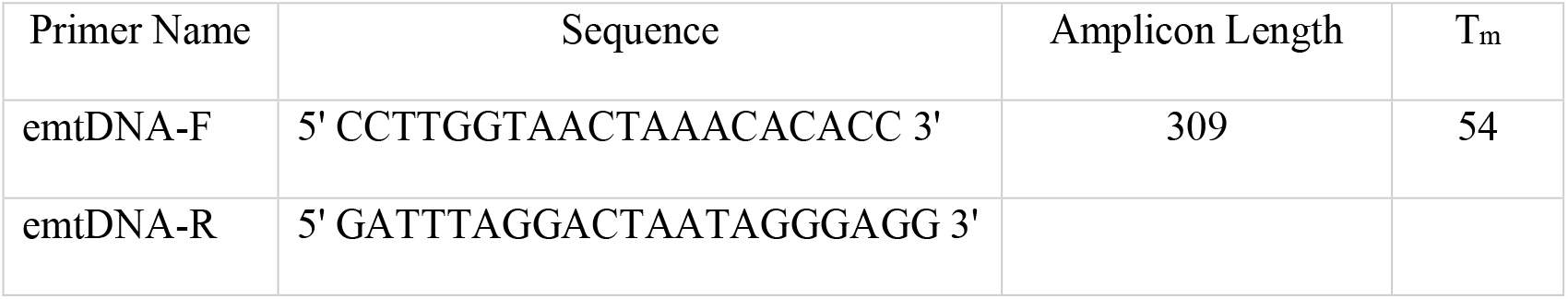
Pāua mitochondrial ATP8-ATP6 eDNA primers for genetic variation.

### 2.4 Mitochondrial eDNA amplicon library preparation and analysis

We used a single-step fusion primer library preparation using our pāua-specific fusion primers for our eDNA samples (Berry et al., 2017; Stat et al., 2017) (Table 3). Prior to any laboratory work, we sterilized all equipment with a 10% bleach solution. We tested eDNA samples to determine which elute (first or second) and which dilution (neat, 1:5, 1:10) would be most suitable (Murray et al., 2015). The second elution with no dilution was found to amplify best (suggesting some slight inhibition) and was used for downstream analyses. No template controls were included with all PCRs. We amplified eDNA samples using 1x SensiFAST SYBR Lo-ROX mix (Meridian Bioscience), 1μM forward fusion primer, 1μM reverse fusion primer, 1μL of template, and 3μL of dH2O on a QuantStudio 3 Real-Time PCR system (Thermo Fisher Scientific) with an initial step at 50°C for 2 minutes, initial denaturing at 95°C for 10 minutes, 40x cycles of 95°C for 15 seconds followed by 60°C for one minute, and then a melt curve of 95°C for 15 seconds, 60°C for one minute, and then an increase in temperature to 95°C with a ramp speed of 0.1°C/second. Sample replicates were test amplified to determine at which cycle they reached plateau, and then again for a final amplification according to which cycle they reached plateau. Amplification was stopped before plateau during the exponential phase of PCR amplification to mitigate PCR bias (Deagle et al., 2009).

PCR products from eDNA were pooled according to amplification cycle and negatives were pooled together to form an eighth pool and spiked into the final library. We cleaned and size-selected these mini-pools according to MagJET NGS Cleanup and Size Selection Kit (Thermo Fisher Scientific) using manufacturer’s instructions and two 85% ethanol washes in a 1:0.9 ratio of PCR product:beads to select for 300bp and above. Cleaned products were measured for concentration with a QuBit dsDNA High Sensitivity Assay Kit on a Qubit 2.0 Flurometer (Invitrogen) and run on a QIAxcel Advanced System (Qiagen) using a DNA High Resolution Kit (Qiagen) with NGS High Resolution Process Profile to determine size and molarity. We measured the final pool with the Qubit 2.0 Flurometer and QIAxcel Advanced System again before submission to Otago Genomics for 2×250 MiSeq Illumina paired-end sequencing using custom read primers FL1-CS1 5’-A+CA+CTG+ACGACATGGTTCTACA-3’ and FL1-CS2 5’-T+AC+GGT+AGCAGAGACTTGGTCT-3’ (M. Bunce, pers. comm.). The “+” indicates linked nucleotides.

After examining read quality with FastQC 0.11.9, we demultiplexed our dual-indexed library using CutAdapt 2.10, excluding indels, with a minimum error rate of 0.15 (maximum error rate of 15%), and minimum size of 100 bp for each read (Andrew, 2020; Martin, 2011). We merged files after renaming with BBMap 38.81, performed a quick quality check with MultiQC 1.9, and size selected for a 309 bp length (Bushnell et al., 2017; Ewels et al., 2016). Primers were trimmed with CutAdapt 2.10 to a minimum length of 268 bp, the amplicon insert size (Martin, 2011). Samples were then filtered with USEARCH 11.0.667, specifying a strict maximum error rate of 0.1 and a minimum of 10 reads per amplicon (Edgar, 2010). Sequences were run through unoise3 to filter real variation and create an OTU table (Edgar, 2016).

### 2.5 Genetic comparisons

We compared haplotypes obtained from tissue and eDNA in Warrington to determine if eDNA could recover true haplotypic variation. Descriptive statistics were used to report mean ± standard deviation (SD) for read counts per replicate in each location. The haplotype percentage across all samples, expressed as individuals per total sampled population (tissue), or haplotype reads per total number of reads (eDNA), was calculated for Warrington. This was further visualized as haplotype percentage for tissue and eDNA.

For tissue, a haplotype network was visualized using a median joining network (Bandelt et al., 1999); in Popart (Leigh & Bryant, 2015). We also compared our tissue haplotypes to previously described ATP6-ATP8 haplotypes of Will et al., (2009). Previous haplotypes from across New Zealand were obtained from Will et al., (2009) and truncated (∼255 bp on the 5’ end) to align with eDNA amplicon haplotypic diversity using Geneious Prime 2021.1.1 (https://www.geneious.com). The six modern tissue haplotypes were also truncated slightly (∼17 bp on the 3’ end) to fit truncated historic haplotypes. The six modern, representative Warrington haplotypes were added to the previous 131 haplotypes to visualize observed and historic diversity using a median-joining network in popart. Importantly, size in the popart visualization does not indicate haplotype abundance; size is a count of how many haplotypes were the same sequence when truncated to eDNA haplotype amplicon size, e. g. how many historic haplotypes matched up to a modern eDNA amplicon.

A haplotype accumulation curve given the haplotypic frequencies seen in the tissue data was modelled with random sampling for 1,000 permutations in R with the s*pecaccum* command in *vegan* to determine how many tissue samples may have been needed to fully acquire all haplotypic variation present at the Warrington site (Oksanen et al., 2020; R Core Team, 2013).

### 2.6 Warrington and Ohau Point eDNA genetic comparison

We compared eDNA haplotypes from Warrington and Ohau Point. Descriptive statistics were used to report mean±SD for Ohau Point read counts. Warrington and Ohau Point haplotype abundance were calculated and visualized by comparing the number of reads for each haplotype to the total number of reads at each site.

## 3. Results

### 3.1 eDNA amplicons obtained

Warrington pāua eDNA amplicons were successfully recovered from 96.3% (n=26/27) of replicates. Most Warrington eDNA samples successfully amplified: nine Warrington samples with three replicates each amplified except for one sample where only two replicates amplified. The Ohau Point sample successfully amplified in all seven replicates (n=7/7, 100%). After quality control, there was a mean±SD of 18,286±15,400 reads per replicate for all locations. When these reads were disaggregated by location, there was a mean of 18,465±16,598 reads (range: 164 – 62,575 reads) per replicate which averaged to 53,344±33,963.64 reads per sample (each sample had 3 replicates, except sample 8 as replicate 8.3 did not yield any reads) for Warrington. Ohau Point replicates yielded a mean of 17,619±10,783 reads (range: 309 – 29,768 reads). Negatives had between zero and two reads per replicate and were not carried through statistical analyses.

Of the nine Warrington eDNA samples recovered, all yielded multiple haplotypes for each read (Figure 4). This held true for the individual replicates as well. Nineteen of 26 replicates had a minor allele frequency that was at least 10% of the major allele frequency for post-processed reads. The most common haplotype overall (Haplotype 1) was found in all replicates except for one (Sample 10, replicate 1). The second most common haplotype overall (Haplotype 2) was found in all replicates, although abundance was not high in all replicates.

**Figure 4.**
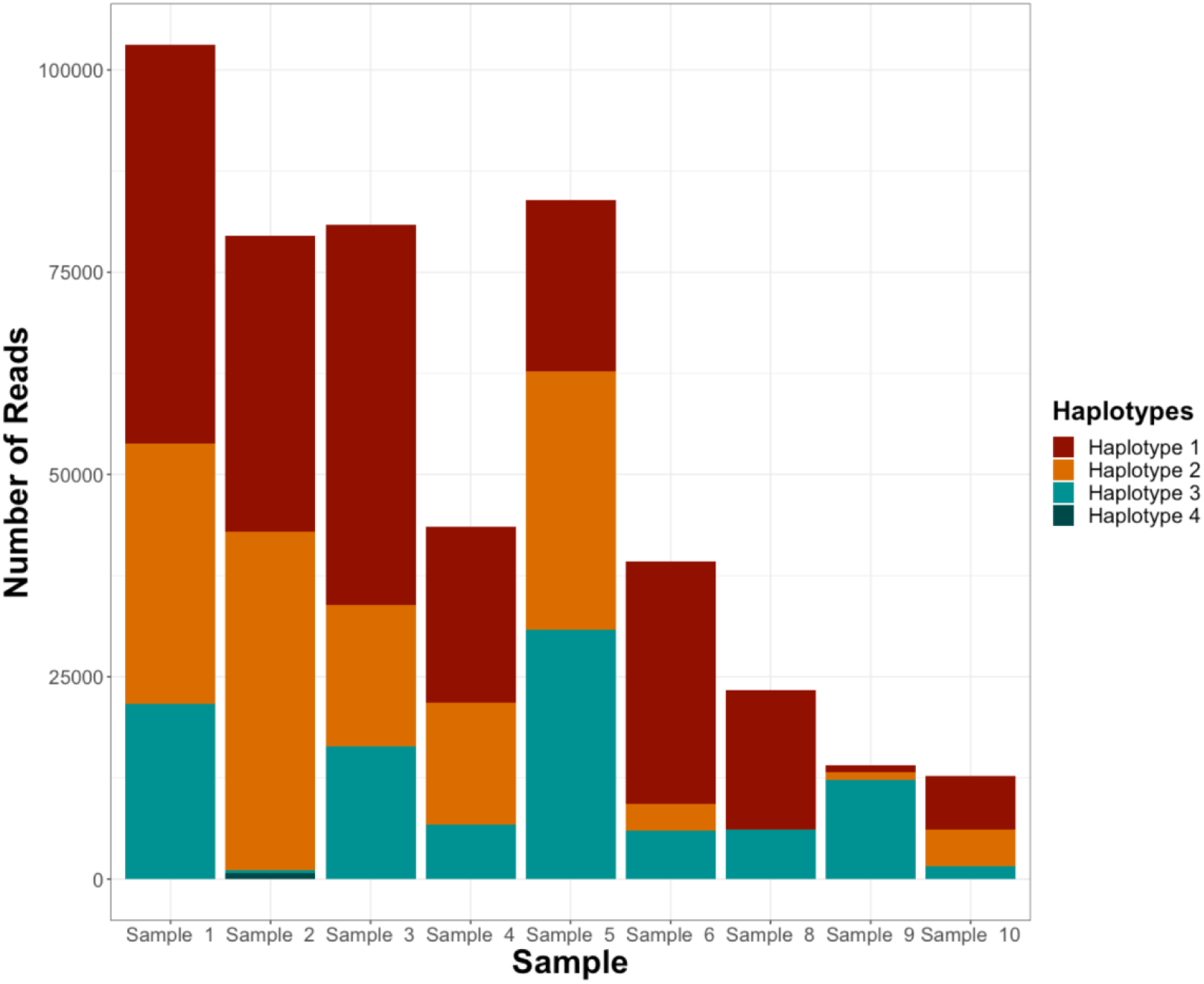
Bar plots showing the number of reads for each haplotype for Warrington by sample. Each sample is an aggregate of three replicates, except for sample eight which is only two replicates due to one having no reads after quality control. Haplotype 4 exists as a very thin yellow line at the bottom of sample two’s read representation.

### 3.2 Tissue and eDNA comparison

Tissue DNA samples (n=71 successfully amplified) from Warrington yielded six mtDNA haplotypes, and eDNA samples reflected the three most abundant haplotypes. Three common tissue haplotypes, haplotypes 1, 2, and 3, were found at frequencies of 35.21%, 50.70%, and 9.86% respectively, and three rare haplotypes, 4, 5, and 6 were found in just one individual each, amounting to 1.41% of the total sampled population for each rare haplotype (Table 4). These tissue haplotypes were visualized with a median-joining haplotype network (Figure 5). The tissue haplotypes were also placed in the greater context of previously identified haplotypes (Figure 6). Because our eDNA amplicon was shorter than PCR amplicons used in previous studies, in some cases multiple previously identified haplotypes matched individual tissue haplotypes obtained in our study. From Warrington eDNA samples, we recovered the three common haplotypes obtained from the tissue samples (haplotypes 1-3), and one rare haplotype—haplotype 4—but did not recover the two other rare haplotypes (5 and 6) (Figure 7). Rare haplotype 4 was only detected with more than five reads in sample two, replicate two (737 reads) (Figure 4). Haplotype 4 occurred in two other samples, but only as 3 reads and as a singleton. Haplotype 4 has three single nucleotide polymorphisms (SNPs) which differentiate it from haplotype 1 (Figure 5). However, when Warrington eDNA samples are bioinformatically analysed alone (dereplication, unoise3 denoising) without the Ohau Point samples, none of the rare haplotypes are recovered, i.e. no rare haplotype 4 detected.

**Table 4.** Table of Warrington tissue and eDNA haplotypes represented as percentages of all total individuals or sequences across all samples.

| Haplotype | Tissue | eDNA |
| --- | --- | --- |
| Haplotype 1 | 35.21% | 47.99% |
| Haplotype 2 | 50.70% | 30.66% |
| Haplotype 3 | 9.86% | 21.20% |
| Haplotype 4 | 1.41% | 0.15% |
| Haplotype 5 | 1.41% | 0.00% |
| Haplotype 6 | 1.41% | 0.00% |

**Figure 6.**
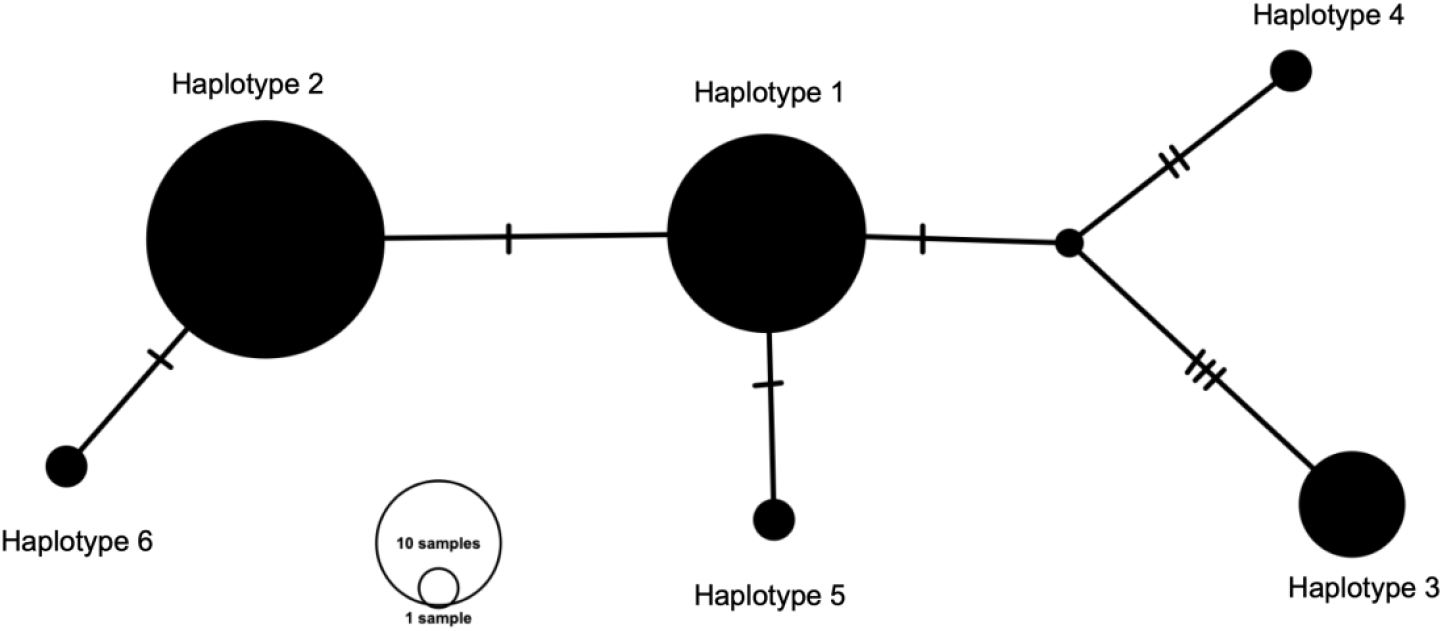
A median-joining network of haplotypes recovered from tissue samples off the coast of Warrington, Otago, New Zealand. Ticks indicate the number of base-pair differences, and size of circle indicates number of individuals with that haplotype.

**Figure 7.**
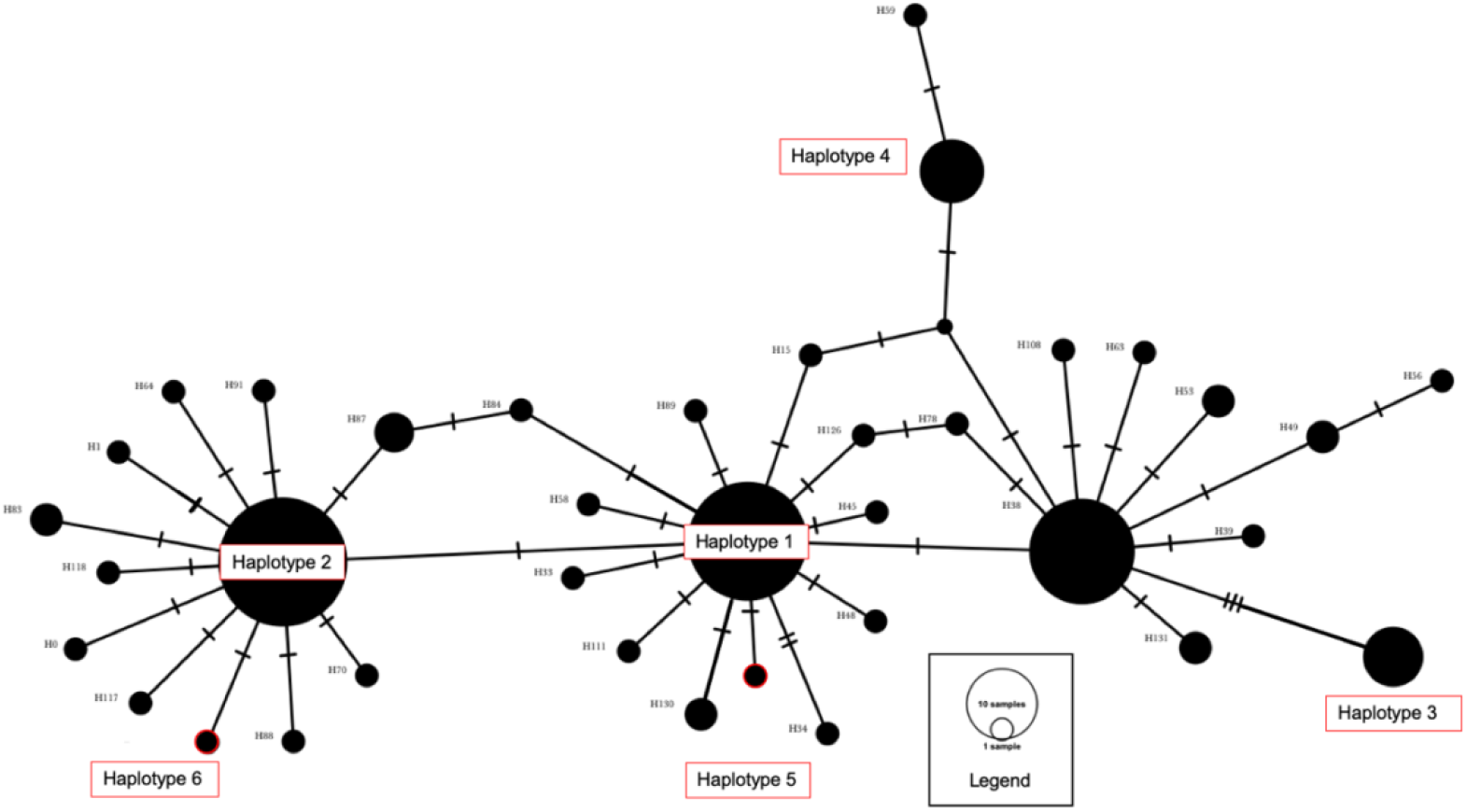
A median-joining network of haplotypes from present-day Warrington tissue representation as well as a count of historical haplotypes from Will et al., 2009. Present-day haplotypes are labelled in red to show where they fit in the haplotype network. One representative (haplotype 1, 2, 3, 4, 5, and 6) from each modern tissue haplotype was used and each of the 131 sequences from Will et al., 2009 were used to build this network. Importantly, size does not indicate haplotype abundance; size is a count of how many haplotypes were the same sequence, e. g. how many historic haplotypes matched up to a modern eDNA amplicon.

A tissue haplotype accumulation curve was produced using one amplicon for each blackfoot pāua individual successfully sampled, yielding a gently sloping curve (Figure 8).

**Figure 8.**
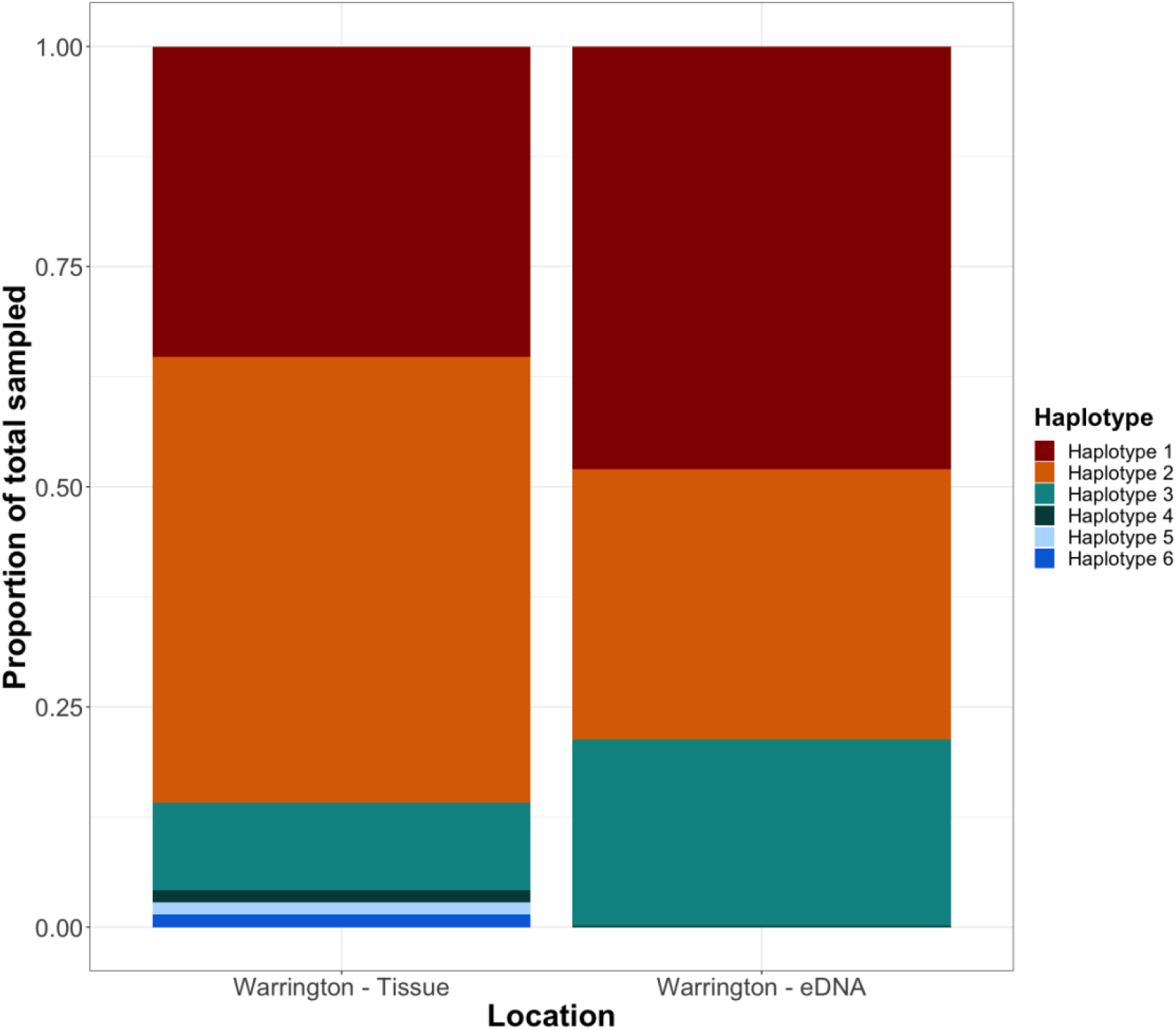
A comparison of haplotypes found in eDNA and tissue. Note haplotypes four through six are rarely found in tissue, and haplotype 4 is only found in eDNA. Haplotype 4 makes up 0.15% of the Warrington eDNA haplotype assemblage and is not visible among other read abundances.

### 3.3 Location comparison of eDNA haplotypes

Four pāua haplotypes were found in both Warrington and Ohau Point eDNA amplicons (1,2, 3, and 4). Haplotype 4 was the rarest haplotype, but was more abundant in the Ohau Point population compared to the Warrington population (Table 5). While the abundances of common haplotypes 1, 2, and 3 differed, all three remained the most common haplotypes found (Figure 9).

**Figure 9.**
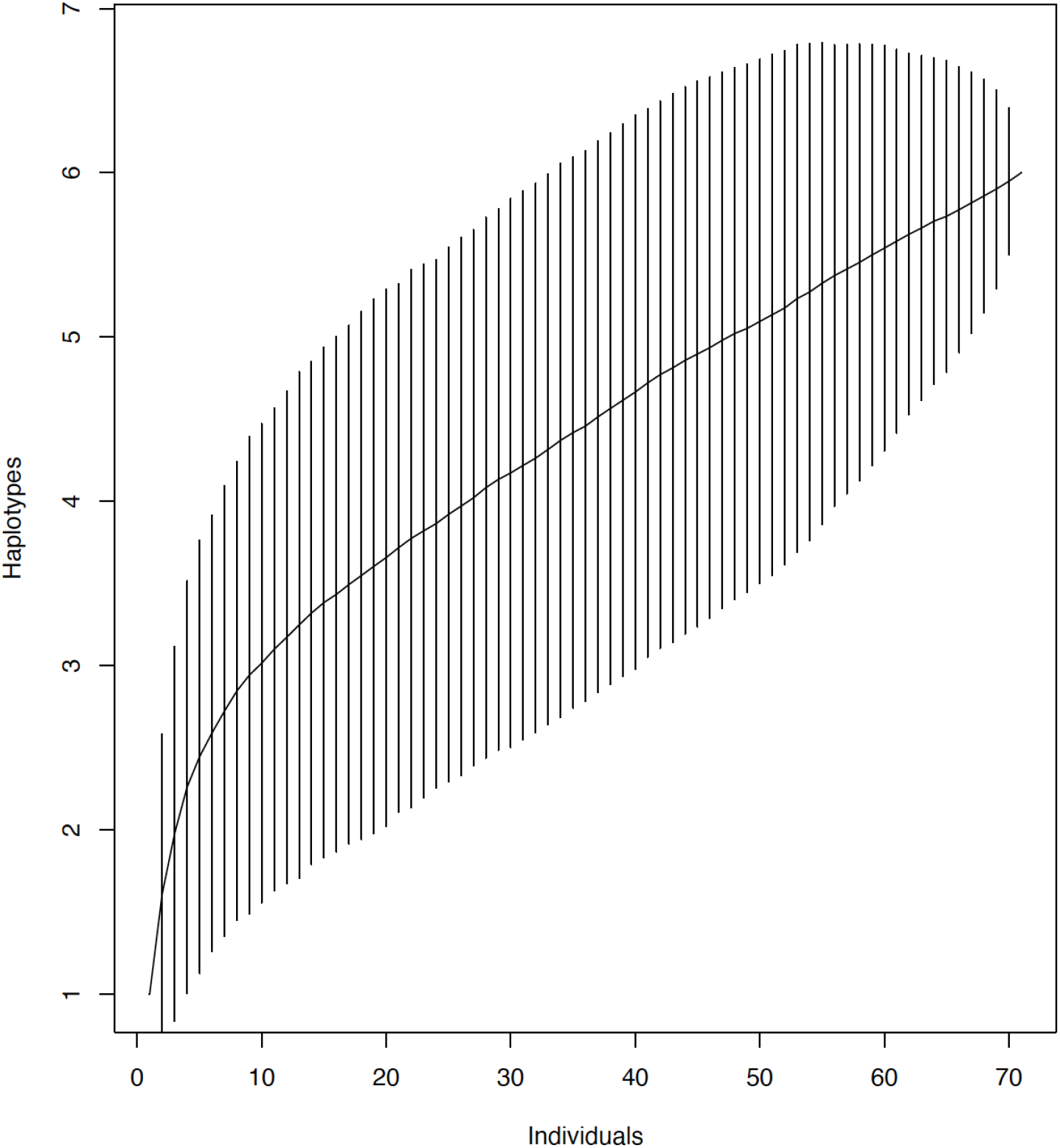
Tissue haplotype accumulation curve based on haplotypes from blackfoot pāua tissue taken at Warrington, Otago. Sequences sampled on the x-axis is equivalent to individuals sampled. Lines indicate confidence interval.

**Figure 10.**
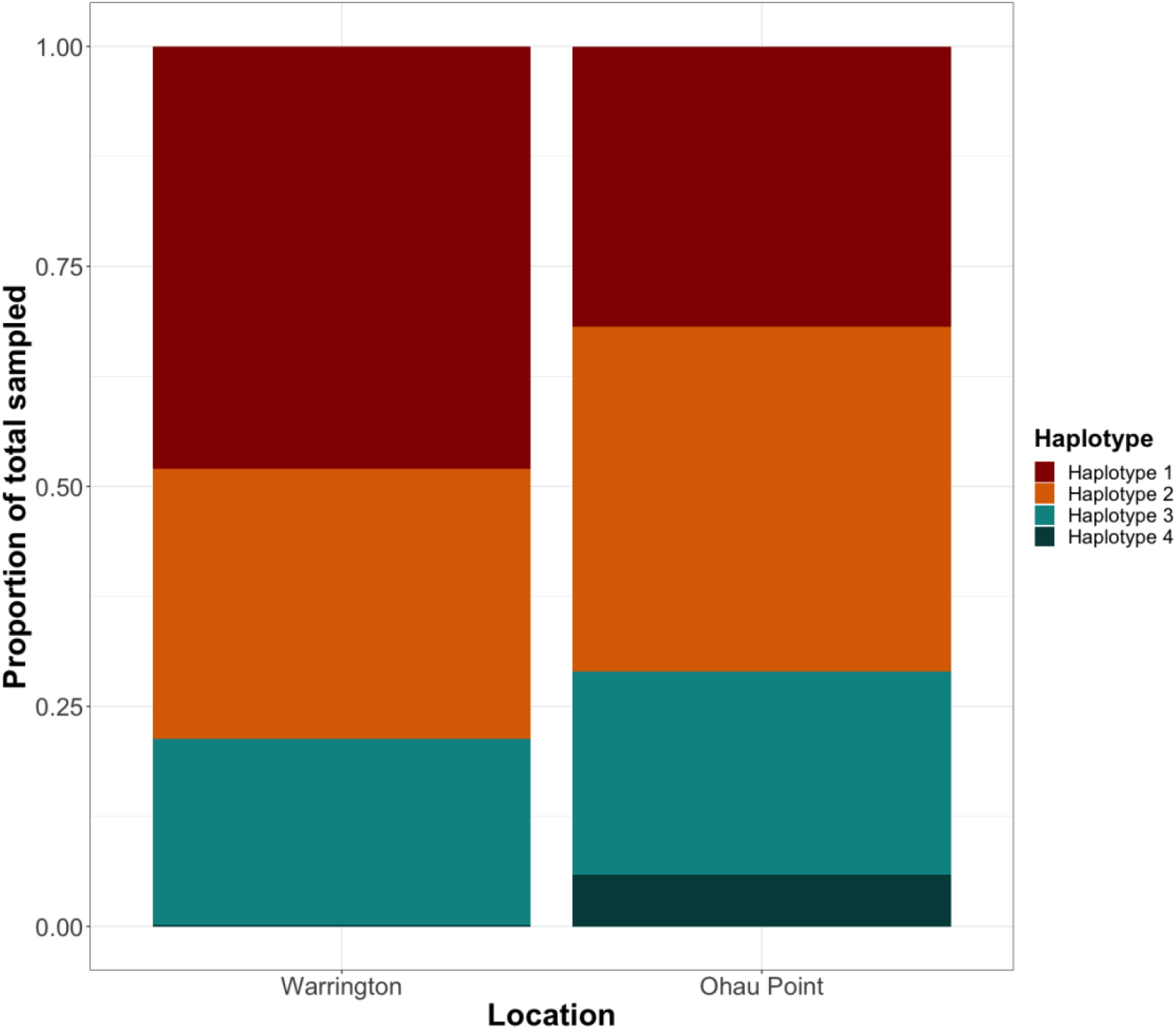
A comparison of Warrington and Ohau Point eDNA haplotypes. Warrington does have a haplotype 4; the number of reads for haplotype 4 was small at Warrington, so it is not easily visually represented.

## 4. Discussion

### 4.1 eDNA identifies common haplotypes found in tissue samples

Here, we show that eDNA can identify the most abundant haplotypic variation present in a population, supporting the idea of eDNA population genetics as a broad-scale monitoring tool. We found three common haplotypes and one rare haplotype for pāua with eDNA methods. Our abundance curve indicates that our eDNA sampling scheme (nine samples, three replicates each) may be comparable to tissue samples (Figure 8). When comparing nine eDNA samples to nine tissue samples, we can see approximately three haplotypes would be identified with tissue sampling. However, Figure 8 indicates increased eDNA sampling effort may be needed to fully capture the rare genetic diversity identified from 70 tissue samples in the Warrington population. This could be because rare haplotypes were more difficult to sample given random chance. Given this tissue haplotype curve, had we taken fewer tissue samples, we would likely have obtained fewer haplotypes (Hale et al., 2012). Most of our Warrington eDNA samples show all three common haplotypes, confirming that eDNA is often a mix of common genetic variation. Other studies have found different results for different organisms such as dolphins, where some samples had only one haplotype when other haplotypes were known to be present in the population. (Parsons et al., 2018). For pāua, this may reflect eDNA from many individuals close together on the same rock. Pāua densities may also play a role in how many haplotypes are recovered because a large, concentrated population size with high mtDNA genetic diversity is not uncommon (Will et al., 2011, 2015). However, specific haplotypic abundances and rare haplotypes still remain mostly obscured, limiting the fine-scale population genetic inferences that can be made from eDNA samples at this time. Our study re-emphasises the importance of being able to ground-truth eDNA population genetic data with comparable traditionally sampled DNA for the species in question.

### 4.2 Haplotypic abundances and rare haplotypes

The eDNA haplotypic abundance we recorded for pāua did not precisely reflect that obtained using tissue samples. Relative haplotype abundances for pāua may be misrepresented due to low concentration eDNA, as seen by common haplotype abundance varying widely between samples. Haplotype 1 was the most abundant in eDNA samples when, according to the tissue samples, haplotype 2 should have been. Surprisingly, haplotype 3, which was only ∼10% of the sampled tissue, was found in abundance in the eDNA samples. Stochastic detection of rare haplotypes, at a tissue frequency of ∼1.5%, suggest our pāua assay was not powerful enough to detect very rare haplotypes (<2% of the population), although it seems a population haplotype frequency of ∼10% is reliably detectable. False negatives due to abundance remain a problem for eDNA population genetics in some species. For instance, it was suspected some rare fish eDNA haplotypes were not identified during migration sampling, although exact tissue-based fish haplotype abundance was unknown at the time (Tsuji et al., 2020). This is in contrast to another study that found rare mussel haplotypes with eDNA, compared to plankton sampling, at <1% of the population (Marshall & Stepien, 2019). Finding the limits of how frequent a haplotype must be in a population to be consistently detected using eDNA methods requires further testing in multiple different species and environments.

The growing field of eDNA population genetics still faces detection challenges for some species like pāua. To use eDNA population genetic information in critical management situations will require avoiding over- or under-estimating genetic diversity, and therefore identifying and filling gaps in our knowledge of how to obtain true diversity is essential (Phillips et al., 2019). In our study, haplotype 4 was likely recovered only because we bioinformatically processed all eDNA samples from Warrington and Ohau Point together. The rare haplotypes 4-6, found in tissue, might occur in extremely low quantities (if at all) in eDNA, but are filtered out because they drop below the threshold for PCR and sequencing noise. Bioinformatically, it is difficult to detect very rare haplotypes due to the fuzzy line between true rare diversity and PCR or sequencing error. Both signals present very few reads in few replicates. Here, haplotype 4 was considered noise if Warrington eDNA samples were analysed alone, but the addition of Ohau Point eDNA samples increased the abundance of rare haplotype 4 in the combined dataset, which enabled us to distinguish this rare eDNA haplotype from the Warrington population. While more Warrington sampling could potentially have found this haplotype, a better use of resources was, arguably, sampling different, genetically variable, locations. Our study supports the importance of sampling at multiple, genetically diverse populations to mitigate haplotype loss associated with eDNA sampling techniques.

### 4.3 Comparisons

While our snapshot of eDNA haplotype diversity reflecting that found in tissue provides one lens to look at the pāua population in Warrington, it is not the full story. Tissue collection may still fail to yield rare haplotypes, just as eDNA methods do. Samples from 2005-2007 indicate blackfoot pāua had a large number of private alleles in two mitochondrial markers (COI and ATP8-ATP6), with four common haplotypes throughout the south-east coast of the South Island of New Zealand (Will, 2009). Our own Warrington tissue haplotypes reflect similar diversity to blackfoot pāua samples taken over a decade ago (Figure 6). However, our short amplicon haplotypes match multiple previous haplotypes because we use a truncated, and slightly shifted, ATP8-ATP6 amplicon (Table 6) (Will, 2009). Importantly, the three major eDNA and tissue haplotypes identified in this study reflect four major previously identified haplotypes identified a decade ago from the south-east coast of the South Island, although no direct comparison exists with Warrington (Will et al., 2011). The reflection of both present-day eDNA and tissue on previous population genetic haplotypes suggests that blackfoot pāua genetic diversity retains common haplotypic variation after a decade, and can be detected with eDNA methods.

**Table 6.** Table of Warrington and Ohau eDNA haplotype representation across all samples. Representation of haplotypes indicated as percentage of total sequences found at each site.

| Haplotype | Warrington | Ohau |
| --- | --- | --- |
| Haplotype 1 | 47.99% | 31.89% |
| Haplotype 2 | 30.66% | 39.16% |
| Haplotype 3 | 21.20% | 23.07% |
| Haplotype 4 | 0.15% | 5.88% |

**Table 7.**
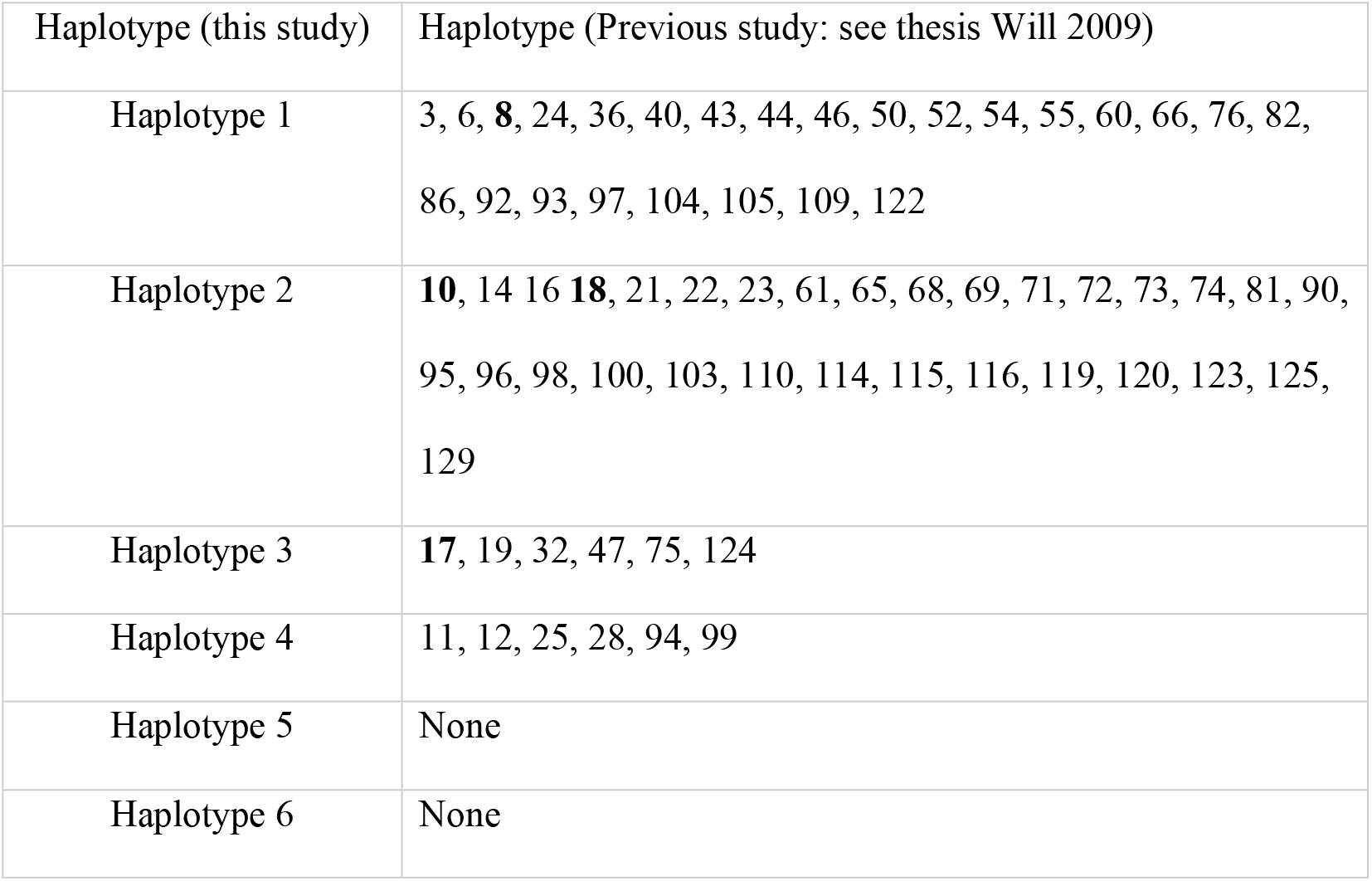
Environmental DNA haplotypes from this study compared with haplotypes from a previous thesis (M. Will 2008). Historical samples were taken 2005-2007 while this study took samples in 2019. Our eDNA haplotypes were represented by many different haplotypes found in previous studies, due to the short nature of our amplicons. The **bold haplotypes (8, 10, 17, 18)** were distinguished in previous studies as abundant (Will 2009, Will et al., 2011).

We find that our pāua eDNA assay amplifies genetic variation from water samples at another location. Our assay detected pāua genetic variation from intertidal water at Ohau Point, in the Kaikōura region, finding the same three common haplotypes detected in Warrington. Tissue-based pāua genetic research in Kaikōura is ongoing (G. Trauzzi, pers. comm.), although current genetic diversity is unknown since the 2016 Kaikōura earthquake. Historic events, such as earthquakes which disturb pāua habitat, need to be considered when choosing eDNA sampling sites, and more ground-truthing for this population is required. The identification of pāua haplotypes via eDNA, despite no identification of live individuals, indicates this assay could be used at locations other than the original Warrington test site.

### 4.4 Recommendations and future directions

For eDNA methods to become a tool in the population genetics toolbox, limitations of intraspecific eDNA techniques need further testing. Without such, meaningful comparisons using eDNA, and building on previously collected samples, cannot be achieved. Based on this study, we have some recommendations for eDNA haplotype sampling.

Most importantly, we recommend ground-truthing eDNA population genetics for each target species, and sampling based on species biology and environment. Biological factors such as hard-shelled animal integument, respiration type and rates, feeding rates (Diggins, 2001; Marshall & Stepien, 2019), and activity levels may have affected organismal eDNA shed and capture (Adams et al., 2019; Allan et al., 2021). For example, grass shrimp (*Palaeomenes sp*.) shed less eDNA than nettle jellies (*Chrysaora sp*.) (Allan et al., 2021). Seasonal effects can also be important, especially those linked to spawning or dispersal. Spawning activities and heightened sexual behaviours have previously led to an increase in aqueous eDNA abundance for some organisms, but this has yet to be tested for pāua eDNA in a field setting (Bayer et al., 2019; Spear et al., 2015; Tillotson et al., 2018; Tsuji & Shibata, 2021). Abiotic and biotic factors also effect the ecology of eDNA itself once shed (Barnes & Turner, 2016). In the dynamic marine environment, eDNA is subject to currents, tides, and salinity (Collins et al., 2018) as well as unique microbial communities (Salter, 2018). It is unknown if these factors affect eDNA haplotypes equally (Berg & Jørgensen, 2006). These factors necessitate species-specific and ecosystem-specific tests.

Secondly, we suggest sampling strategies include multiple locations – especially if there is high genetic variance between populations. Perhaps more importantly, sampling at different locations can help identify rare signal at other sites. Our Ohau Point location allowed us to find the rare haplotype 4 in the Warrington population that otherwise would not have been identified. Sampling across sites may also have the added benefit of highlighting major haplotypes within a population, perhaps even from designing assays that can target local adaptations (Flanagan et al., 2018). More studies comparing the number of sampling locations are needed. Ultimately, as our haplotype accumulation curve shows, more sampling is generally better.

This baseline data highlights a proof-of-concept that real haplotypic variation can be detected in eDNA to some extent, but more work is needed before eDNA can be successfully used to monitor population genetic diversity. Within populations, developing a site-occupancy model for haplotypes may be possible to determine the probability of detecting known but rare variation (Da Silva Neto et al., 2020; Mackenzie et al., 2014). Besides documenting diversity within a population, managers may be interested in gene flow between populations. Adapting an AMOVA using haplotype presence at multiple sites may be of interest to statistically test this, if enough samples have been taken at all target sites (Azarian et al., 2020). Bioinformatic and population genetic tools need to be further developed for eDNA data capturing multiple individuals. Finally, eDNA technology continues to grow, and mitochondrial genomes as well as nuclear molecular markers (microsatellites, SNPs) can now be captured from eDNA (Andres et al., 2021; Jensen et al., 2020; Stepien et al., 2019).

### 4.5 Final remarks

Overall, this research adds to the growing body of literature using eDNA methods for ascertaining haplotypic variation within populations. We suggest that eDNA is currently a broad-scale tool, which readily identifies common haplotypes within a population. We found all common haplotypes and one rare haplotype compared to tissue sampled on the same day at the sample location, confirming eDNA is an effective population genetic sampling tool. Furthermore, we find eDNA haplotypic assays to be robust to application at another location. However, we also note the important lack of detection of rare haplotypic diversity and suggest researchers ground-truth what genetic diversity may be in their own study populations. This specific assay could be a promising route to identifying diversity for other blackfoot pāua populations, but more broadly, eDNA population genetic assays seem extrapolatable to multiple populations in similar habitats once the assay has been confirmed effective for one location. This particular study argues multiple populations are necessary for eDNA population genetic studies, as more varied sampling locations, not depth of sampling at a single site, may be more important to describing genetic diversity and identifying rare haplotypes – especially those rare at one site, but common at another as a result of genetic structure. Finally, this work emphasizes how non-invasive methods can be used to observe genetic diversity at the population level. These non-invasive tools have the potential to reduce harmful interactions between target population and sampler, or may even support research such as behavioural studies which benefit from minimal sampler interference.

## Acknowledgments

Many thanks to members of the Knapp Laboratory for reviewing the manuscript ahead of submission, as well as the Hepburn Laboratory for helping to collect field samples.

